# RNA folding studies inside peptide-rich droplets reveal roles of modified nucleosides at the origin of life

**DOI:** 10.1101/2023.02.27.530264

**Authors:** McCauley O. Meyer, Ryota Yamagami, Saehyun Choi, Christine D. Keating, Philip C. Bevilacqua

## Abstract

Compartmentalization of RNA in biopolymer-rich membraneless organelles is now understood to be pervasive and critical for the function of extant biology and has been proposed as a prebiotically-plausible way to accumulate RNA. However, compartment-RNA interactions that drive encapsulation have the potential to influence RNA structure and function in compartment- and RNA sequence-dependent ways. Herein, we detail Next-Generation Sequencing (NGS) experiments performed for the first time in membraneless compartments called complex coacervates to characterize the fold of many different transfer RNAs (tRNAs) simultaneously under the potentially denaturing conditions of these compartments. Strikingly, we find that natural modifications favor the native fold of tRNAs in these compartments. This suggests that covalent RNA modifications could have played a critical role in metabolic processes at the origin of life.

**One Sentence Summary:** We demonstrate that RNA folds into native secondary and tertiary structures in protocell models and that this is favored by covalent modifications, which is critical for the origins of life.

## Introduction

The RNA World hypothesis states that at the dawn of life RNA may have functioned as both the catalyst and genetic material of life (*1–3*). This has led to great interest in how RNAs and other RNA-like polymers could have been synthesized. Through Miller-Urey experiments, other prebiotic syntheses, and analysis of meteorites, it has been shown that nucleotides/nucleobases could have been present on the early Earth (*4–7*). Also, under certain conditions, a number of amino acids could have been produced prebiotically (*4*, *8, 9*). The solutions resulting from prebiotic chemistries were likely dilute, but encapsulation of these building blocks into a protocell or by adsorption onto mineral surfaces would have greatly increased the concentrations of the prebiotic milieu (*10–16*). This may have set the scene for the synthesis of RNAs that have complex functions akin to modern day riboswitches and ribozymes, which rely on intricate secondary and tertiary structures with complex base pairing at the nucleotide-level to function.

Complex coacervation is a type of liquid-liquid phase separation (LLPS) that has gained plausibility as a robust model for prebiotic (and extant) membraneless compartments (*17*, *18*). Complex coacervates form via a spontaneous associative mechanism whereby polycations and polyanions interact, shedding their counterions, to produce both a dense polymer-rich coacervate phase and a polymer-depleted dilute phase. Frequently in extant biology, intrinsically disordered proteins (IDPs) utilize charged amino acids like aspartate, glutamate, lysine, and arginine to facilitate phase separation, often with nucleic acids (*19–21*). Similarly, polymers of these amino acids can phase separate under a variety of salt and pH conditions (*10*, *22*–*24*). Such coacervates readily compartmentalize RNAs, NTPs, and metal ions (*25*, *26*), all of which would have been essential to functional RNAs on the early Earth. Encapsulation of RNAs inside of peptide-rich coacervate droplets is in many cases strongly favored, however when RNAs are too strongly compartmentalized and interacting with polycations, the RNAs can unfold.

Most investigations into RNA structure and function inside of droplets rely on inferences from ribozyme self-cleavage studies (*26*–*29*), and it has remained challenging to study RNA structure at the nucleotide-level inside of coacervates (*10*, *22*, *23*). Although studies of RNA catalysis provide evidence for adoption of the folded state (*26*–*30*), they are limited as they provide no structural insights at the single-nucleotide level, and what is observed can be due to a very small population of properly folded ribozymes (*31*). Other studies have been performed to determine which RNAs extracted from extant cells partition into various condensates (*32*). Though these studies inform us on the sequence specificity of uptake by condensates, they do not speak to the secondary and tertiary structures adopted by those RNAs within the compartments. Multiple techniques have been recently developed to study RNA structure at the single-nucleotide level for both *in vitro* and *in vivo* applications, some from our own laboratory, opening the door to such investigations (*33*–*38*). For example, recent work has begun to address the lack of high resolution RNA structural studies inside of membraneless compartments (*10*, *26*, *28*, *39*–*41*). It has been demonstrated that biocondensates can alter simple RNA duplex thermodynamics (*22*, *42*), but their impact on RNAs with complex secondary and tertiary structures (typical of functional RNAs), remains largely unknown.

To understand the effects of encapsulation on RNA structure, we sought to study a well-characterized RNA that adopts a complex secondary and tertiary structure. We chose to study a series of post-transcriptionally-covalently-modified and unmodified tRNAs because they have well-defined secondary and tertiary structures with precise positioning of nucleotides analogous to those of ribozymes, and are thought to be an ancient RNA because of their ubiquity among the domains of life (*43–45*). tRNAs also have diverse functions such as supplying various fragments that have key roles in regulating gene expression in extant biology (*46*). Additionally, tRNAs are the most heavily modified and among the most deeply studied RNAs (*47*–*52*). In extant biology, these modifications provide protein binding sites, epigenetic control of expression, roles in the stabilization of native RNA folds, and protect against degradation (*53*, *54*). For example, 2’O methylation increases RNA backbone stability by preventing nucleophilic attack by the 2’-OH, and modifications to the nucleobase can either increase folding stability, as in the case of pseudouridine (*55*), or decrease it, as in the case of dihydrouridine (*56*, *57*). Additionally, modifications on the Watson-Crick face of the nucleobases can prevent base pairing, thereby preventing misfolds (*58*, *59*). Although the above studies have shed light on the effects of modifications on RNA folding, it is unclear what their consequences are on RNA folding inside of protocell models such as the peptide-rich droplets studied here.

Herein, we use confocal microscopy to identify conditions under which complex coacervates form and characterize their physical properties as tRNA-containers by fluorescent microscopy and Fluorescence Recovery After Photobleaching (FRAP). Then, we perform In-Line Probing (ILP) on a single tRNA to define which experimental conditions support nucleotide-specific native secondary and tertiary structure formation. We then take a subset of these conditions and perform our newly described tRNA-Structure-Seq approach (*60*) on ^~^40 naturally modified tRNAs from *E. coli* and their unmodified T7 transcript counterparts inside several different coacervate conditions. These studies reveal for the first time that natural modifications strongly favor RNA folding inside of coacervates, which has profound implications on origins of life scenarios.

## Results

### Oligolysine-oligoaspartate coacervates have similar physical properties under diverse conditions

RNAs in modern cells as well as those that have been envisioned in RNA Worlds have intricate functionally important secondary and tertiary structures. In this study we sought to understand at the nucleotide level whether functional RNAs could fold robustly into their tertiary structures inside of membraneless compartments. Our earlier studies showed that RNA can form simple duplexes and that ribozymes can have modest activity in certain coacervates particularly those with excess polyanion (*22*, *29*). However, it was unclear whether encapsulation would be compatible with robust folding of RNAs with complex secondary and tertiary structures, which are present in all known functional RNAs. Specifically, ILP studies under a limited set of conditions showed that long-range secondary and tertiary interactions did not form inside similar coacervates (*10*). To elucidate conditions where robust RNA folding might be possible, we designed an experimental pipeline amenable to studying RNA structure in peptide-rich droplets (Figure 1) and focused on three experimental variables: polyion (+/-) charge ratio, Mg^2+^ concentration, and polymer length.

**Fig. 1:**
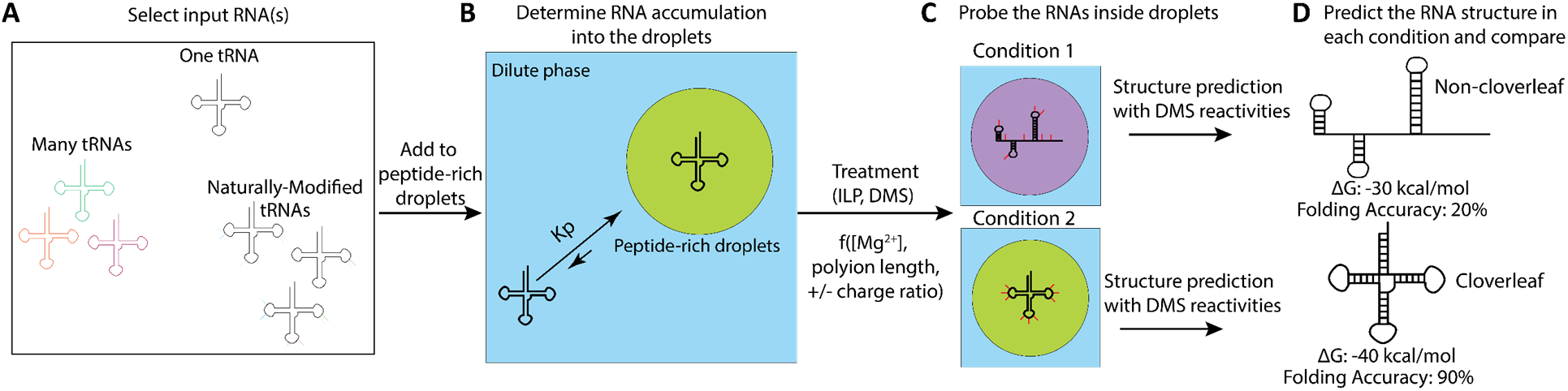
Structure probing of tRNA inside of peptide-rich droplets. **A** As inputs to the droplets one can study one tRNA by both ILP and tRNA structure-seq, many tRNAs by tRNA structure-seq, and even many naturally modified tRNAs by tRNA structure-seq. **B** The level of accumulation into the peptide-rich droplets should be evaluated and then **C** the tRNA(s) can be studied by both ILP and tRNA structure-seq. From there with DMS reactivities derived from tRNA structure-seq experiments, **D** the structure of the RNAs in the condition can be predicted with RNAstructure or other RNA structure prediction software.

Coacervates comprised of either pairs of 10mers or 30mers of oligolysine and oligoaspartate homopolymers were studied, and polyion +/- charge ratios of 1:1, 1:2 and 1:3 were tested (10 mM + charge: and either 10, 20, or 30 mM – charge). Excess polyanion was tested because previous studies from our labs showed enhanced ribozyme catalysis under these conditions (*29*). Folding of RNA tertiary structure in dilute solution is often favored by Mg^2+^. We therefore chose Mg^2+^ concentrations of 0.5 mM and 10 mM to explore RNA folding conditions in membraneless compartments, which reflect typical *in vivo* and *in vitro* concentrations of Mg^2+^ (*61*). Experimental conditions were identified wherein coacervates would form according to published data (*10*) and were supported by light microscopy (Figure S1). Next, we used both fluorescent and radioactive methodologies to test whether a model tRNA (Yphe) was accumulated into the droplets under these conditions (Figure 2A-B, Figure S2, Tables S1-S3). Indeed, Yphe strongly accumulated in the droplets with (90% or greater RNA in peptide-rich phase), achieving up to 230X increases in concentration (Supplemental Discussion 1, Figure S2).

**Fig. 2:**
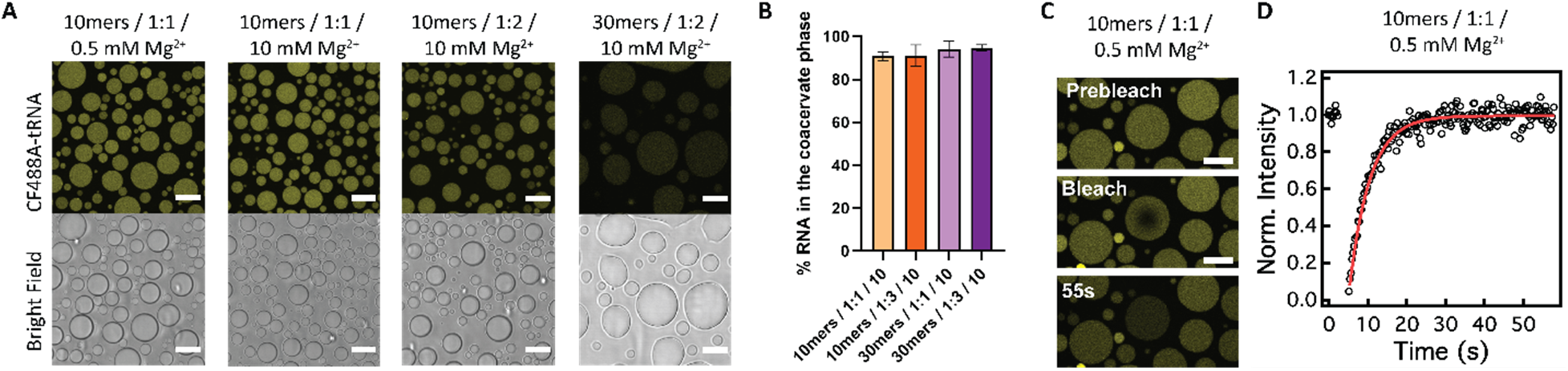
OligoLys-oligoAsp droplets have similar physical properties as judged by fluorescent and radioactive partitioning experiments and FRAP. **A** Fluorescence and brightfield images of CF488A-tRNA inside peptide-rich droplets. Scale bars = 10 μm. **B** Radioactive accumulation of Yphe into peptide-rich droplets. All pairwise comparisons were made with the Kruskal-Wallis test and Dunn’s multiple comparisons test with α = 0.05, but none were significant (Table S2). **C** Fluorescence images of intradroplet bleaching and recovery of CF488A-tRNA in 10mers / 1:1 charge ratio / 0.5 mM Mg^2+^. Scale bars = 10 μm. **D** Representative FRAP curve and fit from the images in **C**.

To test how liquid-like the tRNA containing droplets were, we performed intra-droplet Fluorescent Recovery After Photobleaching (FRAP) with 3’-end-CF488A-labeled Yphe (Figure 2C, D, Supplemental Equations 1, and 2). In the 10mer coacervates, recovery after photobleaching was similar regardless of Mg^2+^ concentration or charge ratio and relatively rapid, with an average half-life of 3.6 ± 1 sec. In 30mer coacervates, the half-life of recovery was somewhat longer, 7.0 ± 1 sec (Figure S3A, Table S4). Like the half-time of recovery, the apparent diffusion coefficient was similar amongst the 10mer coacervates (Figure S3B, Table S5, Supplemental Equation 3). Again, the 30mer coacervates had slightly slower diffusion. Upon establishing partitioning of tRNA and droplet fluidity, we moved to examine the fold of Yphe at the single nucleotide level under those conditions.

### In-Line Probing of an unmodified tRNA identifies conditions that favor native folding

To isolate the individual impacts of +/- charge ratio (1:1 vs 1:2), Mg^2+^ concentration (0.5 mM vs 10 mM), and polymer length (10mers vs 30mers) on the fold of Yphe, an unmodified tRNA, we designed a conceptual cube where a single variable changes along each edge (Figure 3A). Each face of the cube held one condition constant, such as n=10 for the polyion length (Figure 3B). With this formalism, we performed In-Line Probing (ILP) experiments with the 5’-end ^32^P-labeled T7 transcript of Yphe in the conditions defined on the face in Figure 3B, where circle size depicts the relative quality of folding as judged by the reactivity of the variable loop (Figure 3C, Supplemental Methods, Table S6).

**Fig. 3:**
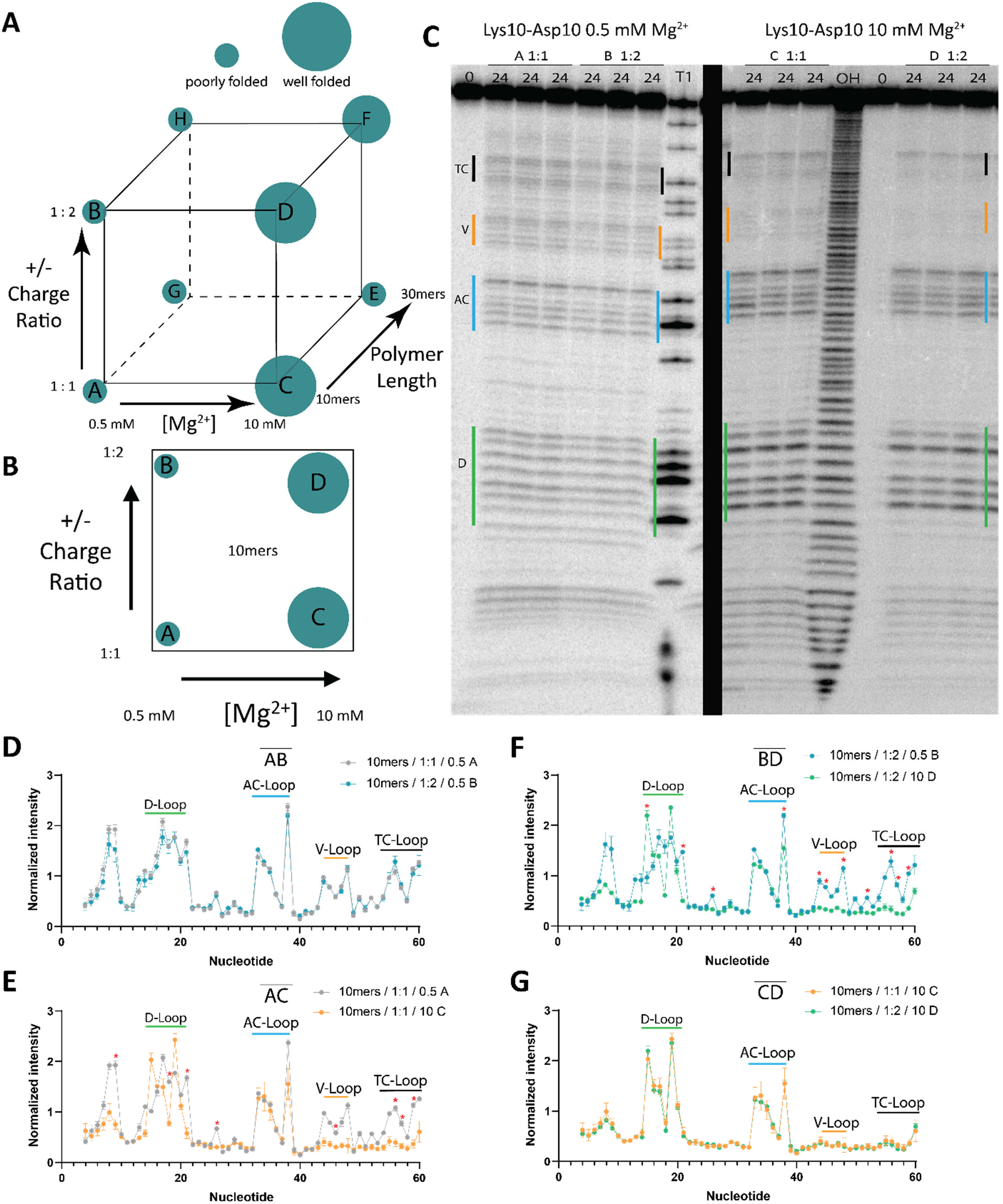
Isolating the effects of changing multiple parameters on In-line probing (ILP) of Yphe. **A** Conceptual cube with each vertex assigned a condition, where edges indicate a change in either [Mg^2+^], polyion length, or +/- charge ratio. Small circles indicate poor folding and large circles indicate better folding **B** Face ABDC of the cube, where polymer length n=10 is constant. Teal circles indicate how well Yphe folded. **C** Gel image of Yphe ILP in Conditions A, B, C and D. **D-G** Overlay line graphs of ILP reactivities from conditions **D** edge AB, **E** edge AC, **F** edge BD, and **G** edge CD. Line graphs represent the data from three independent experiments per condition and error bars show the standard deviation of the normalized intensity values at each point. Intensities were normalized to the average intensity of nucleotides 34-36. Red asterisks indicate significant p-values as judged by multiple t-tests, one at each point with α = 0.05 and the Holm-Sidak method to correct for multiple comparisons (**Table S7**).

Our previous ILP studies showed that 0.5 mM Mg^2+^ supported robust folding of Yphe in dilute solution, but not in similar peptide-rich droplets (*10*). In 10mer droplets, at 0.5 mM Mg^2+^ Yphe was poorly folded in both 1:1 and 1:2 +/- charge ratios corroborating the previous results (edge AB, Figure 3D, Table S7). At both 1:1 and 1:2 charge ratios, Yphe was better folded at 10 mM than 0.5 mM Mg^2+^ (edges AC and BD, Figure 3E-G, Table S7). To summarize, in 10mer coacervates, increasing the Mg^2+^ concentration significantly increased the ability of Yphe to fold natively, and here the +/- charge ratio did not affect the fold.

With this knowledge, we turned to the other five faces of the cube to make additional comparisons of tRNA folding in the coacervates (see Supplementary Discussion 2, Figures S4-S12, and Tables S8-16). In contrast to 10mer coacervates where 10 mM Mg^2+^ was sufficient for folding, in 30mer coacervates Yphe required both 1:2 +/- charge ratio and 10 mM Mg^2+^ to fold well (Figure S8). At 0.5 mM Mg^2+^, Yphe did not fold well under any condition, although it folded somewhat better in 10mer than 30mer coacervates. (Figure S9-10). At 10mM Mg^2+^ and 1:1 charge ratio, Yphe folded better in 10mer than 30mer coacervates but folded equally well in both polymer lengths at 1:2 charge ratio (Figure S11-12). Overall, native folding of Yphe was favored by higher [Mg^2+^], shorter polymer lengths, and excess polyanion. We interpret these observations as being due to the following effects: higher [Mg^2+^] provides greater charge neutralization, shorter polymers have fewer possible modes of interaction that can disrupt the fold of the RNA, and excess polyanion likely displaces denaturing RNA: polycation interactions. Notably, the ability of Yphe to adopt its proper fold inside oligoLys/oligoAsp droplets depends on the interplay of these factors.

### Development and application of tRNA structure-seq in peptide-rich droplets

To gain greater insight into how multiple tRNAs fold simultaneously, we next turned our attention to chemical probing with readout by NGS. This was important because ILP cannot handle large numbers of conditions and sequences, and it currently does not provide parameters to help predict the fold of RNA. In this section, we describe the application of NGS to a single tRNA in droplets for the first time, while in the next section, we expand this method to study ^~^40 different tRNAs in droplets. We recently developed tRNA structure-seq, which applies dimethyl sulfate mutational profiling (DMS-MaP-Seq) to elucidate the secondary structures of all of the tRNAs in an organism.(*60*) To select conditions, we returned to the conceptual cube from the ILP studies (Figure 3A). From that analysis, we chose to perform tRNA-Structure-Seq studies under the following four representative sets of conditions: Condition A: 10mers/1:1/0.5, Condition C: 10mers/1:1/10, Condition D: 10mers/1:2/10, and Condition F: 30mers/1:2/10 (Lys-Asp length/ +/- charge ratio/ [Mg^2+^] in mM). These experiments required higher concentrations of RNA than ILP (see Materials and Methods). As such, we performed radioactive RNA accumulation experiments at the higher RNA concentration, and Yphe was strongly accumulated into the coacervates (again, >90 % of Yphe was encapsulated under all conditions, Figure S13, Table S17). We then prepared tRNA-Structure-Seq libraries (see Materials and Methods) under Conditions A, C, D and F. Analysis by ShapeMapper2 (*36*) provided mutational profiles of Yphe in each of these four conditions, with each experiment performed in triplicate. DMS reactivities from the replicate libraries were well correlated (R^2^ ≥ 0.89) (Figure S14, Table S18), multiple mutations per read in the +DMS conditions were detected (Figure S15), and characteristic mutations at residues A, C, and G were reliably identified (Figure S16, Table S19). Reactivities at A and C report on the accessibility of the Watson-Crick faces, which provide empirical restraints for RNA structure prediction with folding software such as RNAstructure (*62*, *63*).

When the DMS reactivities from Condition A (10mers/1:1/0.5) were used to restrain prediction of the structure, Yphe folded into a rod-shape rather than its characteristic cloverleaf structure, with a folding accuracy of just 26.7% (Figure 4B, Table 1). Remarkably, increasing the Mg^2+^ concentration to 10 mM in Conditions C, D, and F, led Yphe to fold into its cloverleaf structure rather than a rod-shape (Figure 4C-E). This led to a significantly improved folding accuracy of 97.6% in these three conditions (Table 1) and lowered the apparent free energy ΔG_37_ of the predicted structures from −24.3 to −27.9 to - 34.6 kcal/mol across these three conditions, with the latter two being in excess polyanion. This quantitative result matches the trends from ILP experiments above and provides a mechanistic explanation for findings from Poudyal et al. where excess polyanions led to enhanced ribozyme activity (*29*).

**Table 1:** ΔG_37_, Sensitivity, PPV, and Accuracy values for folding Yphe with various pseudo-free energy parameters, for the predicted structures in **Figure 4 A-D**.

| Condition | $\Delta G_{37}$<br>(kcal/mol) | Sensitivity | PPV | Accuracy |
| --- | --- | --- | --- | --- |
| *in silico* | -22.7 | 95.2% | 100% | 97.6% |
| 10mers / 1:1 / 0.5 (A) | -26.5 | 28.6% | 25% | 26.7% |
| 10mers / 1:1 / 10 (C) | -24.3 | 95.2% | 100% | 97.6% |
| 10mers / 1:2 / 10 (D) | -27.9 | 95.2% | 100% | 97.6% |
| 30mers / 1:2 / 10 (F) | -34.6 | 95.2% | 100% | 97.6% |

**Fig. 4:**
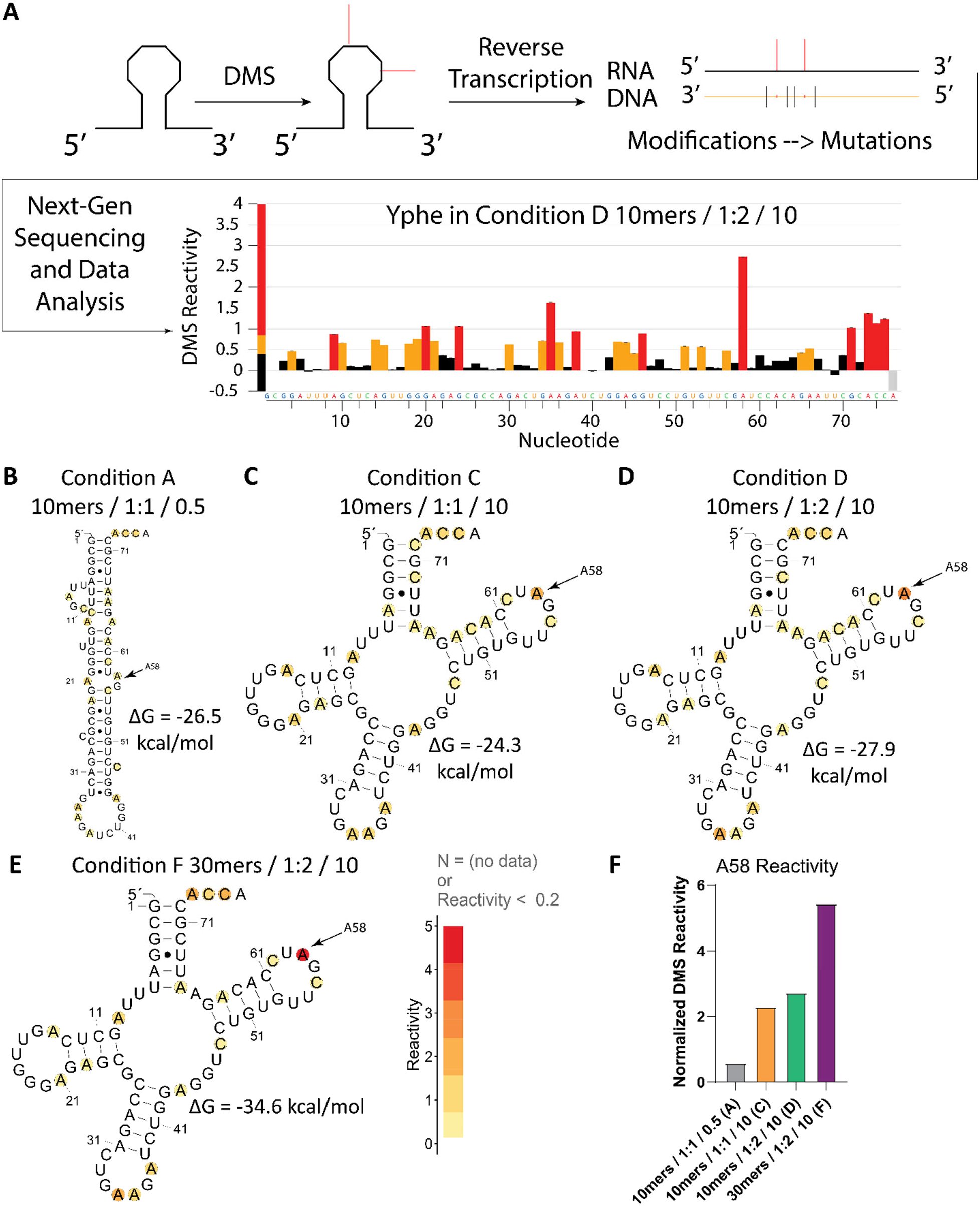
tRNA structure-seq in droplets identifies conditions for robust native folding of tRNA. **A** Scheme of DMS-MaP reaction and data analysis to get the DMS mutation rate at each nucleotide. **B-E** Predicted structures of Yphe in **B** Condition A (10mers / 1:1 / 0.5), **C** Condition C (10mers / 1:1 / 10), **D** Condition D (10mers / 1:2 / 10), and **E** Condition F (30mers / 1:2 / 10). **F** Normalized DMS reactivity of nucleotide A58 under each condition. Bars represent the ShapeMapper2 normalized reactivity of three combined replicates and do not have error bars because they were combined before normalization. Structures in B-E were generated with R2easyR (*88*) and R2R (*89*).

In all known tRNAs, nucleotide A58 forms a reverse Hoogsteen base pair with U54, which orients A58 into solution allowing it to be hypermodified by DMS, if it is not already naturally methylated (*60*, *64*). Indeed, hypermodifcation of A58 by DMS is consistent with formation of the TΨC stem-loop (*65*). Going across all four conditions (A, C, D, F), there was a sharp increase in the DMS reactivity of A58 (Figure 4F). This trend supports the enhanced folding of the TΨC stem-loop, which is present in both tRNA secondary and tertiary structures, across these four conditions. Having verified that we could perform tRNA structure-seq on a single tRNA inside of the coacervates, we next moved to study multiple tRNAs in coacervates simultaneously.

### tRNA structure-seq in peptide-rich droplets reveals a role of natural modifications in limiting tRNA misfolds

After successfully studying a single tRNA at the single nucleotide level in droplets by the tRNA structure-seq (*60*) method, we applied it to simultaneously investigate the effects of encapsulation in droplets on ^~^40 unmodified T7 transcripts (T7) of *E. coli* tRNAs and their naturally modified (NM) counterparts. This provides an outstanding system to investigate effects of modification on RNA folding as *E. coli* tRNAs have well characterized nucleotide modifications that vary in size, shape, hydrophobicity, and ability to basepair (*66*). While much is understood about enzymes that install these modifications (*67*–*69*), little is understood about the synergistic effect of these modifications on the folding of their tRNAs, particularly in droplets (*52*, *70*). Understanding the effects of natural modifications on folding is significant to origins of life scenarios where modified nucleosides may have been prevalent (*51*, *54*, *71*).

Experiments were performed as in the previous section in Conditions A, D, and F. Analysis of the data by ShapeMapper2 (*36*) provided mutational profiles for the sets of T7 and NM tRNAs under each of these three conditions. The DMS reactivities of replicate libraries were well correlated (R^2^ ≥ 0.91) (Figure S17, Table S20). In the +DMS conditions, there was a higher proportion of multiple mutations per read than in -DMS conditions (Figure S18), and mutations at residues A, C, and G could be detected reliably (Figure S19, Table S21). In total, we had sufficient coverage under all three conditions for 39 of the 49 T7 tRNAs, and 45 of the 49 NM tRNAs (Table S22).

We assessed the apparent ΔG_37_ of each tRNA by folding them using the DMS reactivities of A and C as pseudo-free energy restraints. The NM tRNAs had significantly lower mean apparent free energies ΔG_37_ than the T7 tRNAs, by 7-10 kcal/mol, across all three conditions, consistent with the NM tRNAs folding more robustly (Figure 5A, Tables S23-24). Moreover, the mean reactivity at A58 was significantly greater for the NM tRNAs than the T7 tRNAs across all three conditions, especially under the most stringent folding condition (A) (Figure 5B; Table S25), strengthening the above conclusion. The nucleotide level resolution of the data can be displayed on heatmaps (Figure 5C,D, Figures S20,S21). Two features of the DMS reactivity at each nucleotide stand out with respect to the robust folding of the NM tRNAs. The reactivity at A58 and at the terminal CCA nucleotides are noticeably higher for NM tRNAs than T7 tRNAs. These two trends were also present in Conditions D (10mers/1:2/10) and F (30mers/1:2/10) (Figures S20-21), although they were clearest under Condition A (10mers/1:1/0.5), again suggesting that RNA modifications have the greatest impact under the most stringent folding condition (A).

**Fig. 5:**
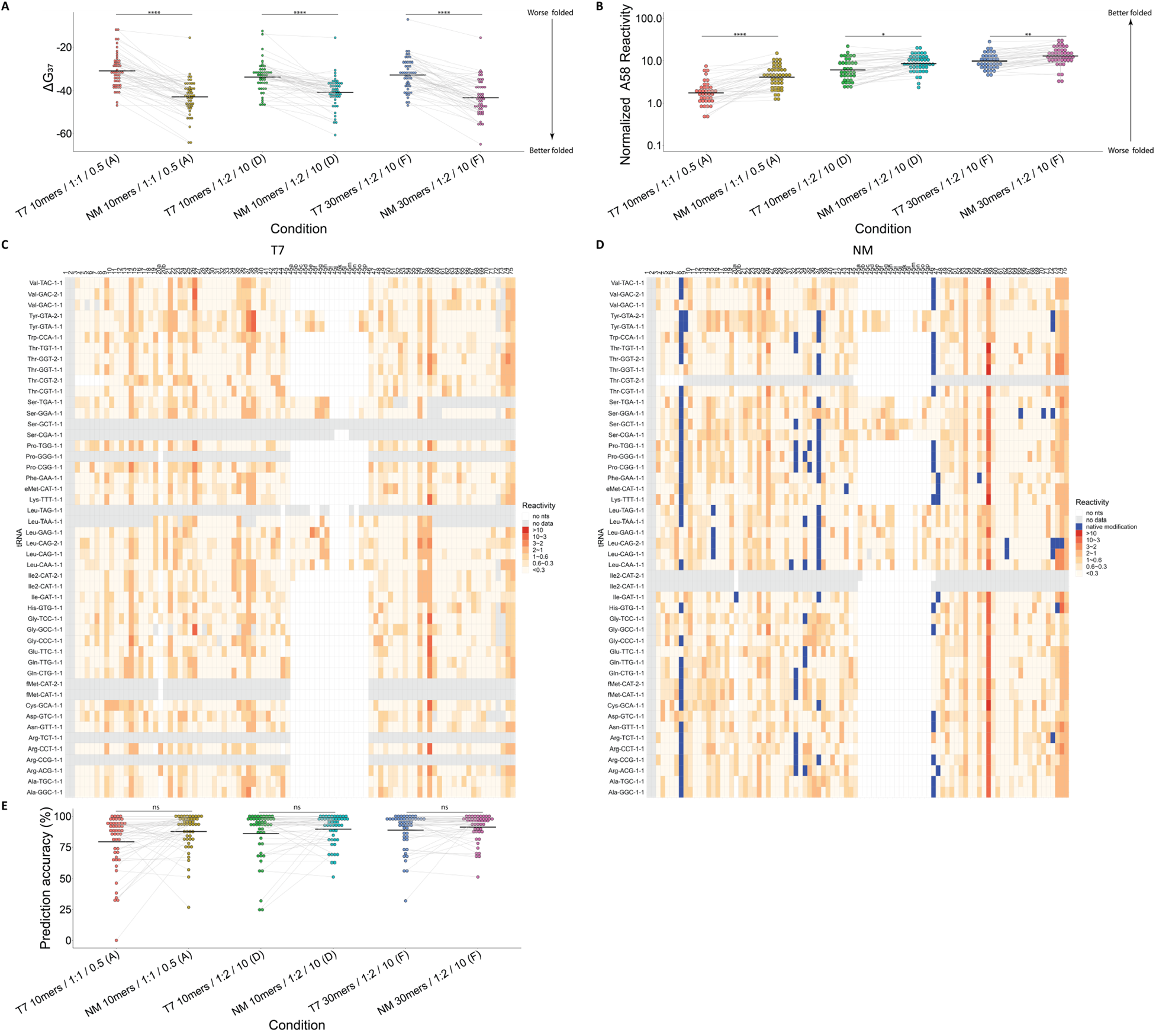
Natural modifications drive native folding of tRNAs in droplets. **A** Apparent ΔG_37_ of each T7 and NM *E. coli* tRNA in each condition. **B** Normalized DMS reactivity at position A58 for each T7 and NM *E. coli* tRNA under each condition. **C/D** Heatmaps of DMS reactivity at each nucleotide for each T7 **C** and NM **D** *E. coli* tRNA in Condition A (10mers / 1:1 / 0.5). **E** Prediction accuracies of T7 and NM *E. coli* tRNAs in each condition. For **A**, **B**, and **E** significance was evaluated by Dunnett’s T3 multiple comparisons test. Level of significance is indicated by “ns” for p-values greater than 0.05 “*” for p-values less than 0.05, “**” for p-values less than 0.01, “***” for p-values less than 0.001, and “****” for p-values less than 0.0001. Significant p-values are shown between only T7 and NM *E. coli* tRNAs in the same conditions, all other comparisons are shown in **Tables S24-25**, and **S31.**

Given the favorable influence of natural modifications on tRNA folding in peptide-rich droplets, we wanted to test whether they could improve the accuracy of tRNA structure prediction, where accuracy is a measure of how similar the in-droplet DMS-restrained predicted structures are to the known structures. To accomplish this, the structures of the 49 tRNAs were predicted in dilute solution (*in silico*) with RNAstructure (*63*) and compared to their known structures from tRNADB (*72*) using the scorer function of RNAstructure. For these predictions, the mean accuracy (an approximation of the Matthew’s Correlation Coefficient (MCC), see Supplemental Equation 4) was modest at 79.1% (Table S26). Next, the T7 and NM tRNAs were folded using their DMS reactivities from each droplet condition as pseudo-free energy restraints (Figures S22-S27, Tables S27-S30). Upon comparison to their known structures from tRNADB, both T7 and NM tRNAs folding accuracy improved as [Mg^2+^], charge ratio, and polymer length increased (Table 2). Strikingly, under all three conditions, the NM tRNAs had higher mean accuracies than their T7 transcript counterparts even though the difference in the distributions was not significant (Figure 5E, Table S31). Once again, RNA modifications had the most impact on the mean accuracy under stringent folding condition (A).

**Table 2:** Average Sensitivity, PPV, and Accuracy values for *in silico* predictions, T7, and NM E. coli tRNAs. (Folded structures and tables containing all of the individual values for sensitivity, PPV, and accuracy can be found in **Figures S22-S27** and **Tables S26-30**).

| Condition | Sensitivity | PPV | Accuracy |
| --- | --- | --- | --- |
| *in silico* | 81.6 | 76.8 | 79.1 |
| T7 10mers / 1:1 / 0.5 (A) | 76.4 | 82.6 | 79.3 |
| T7 10mers / 1:2 / 10 (D) | 84.2 | 88.0 | 86.0 |
| T7 30mers / 1:2 / 10 (F) | 86.9 | 90.8 | 88.7 |
| NM 10mers / 1:1 / 0.5 (A) | 88.3 | 87.1 | 87.6 |
| NM 10mers / 1:2 / 10 (D) | 89.8 | 89.5 | 89.5 |
| NM 30mers / 1:2 / 10 (F) | 91.0 | 91.6 | 91.2 |

Prokaryotic and eukaryotic cells have varying intracellular Mg^2+^ concentrations, and Mg^2+^ ions are essential for RNAs to fold into their tertiary structures (*70*, *73–78*). We therefore analyzed the sensitivity of the various T7 and NM tRNAs to Mg^2+^ concentration between 0.5 and 10 mM, which represent typical *in vivo* and *in vitro* Mg^2+^ concentrations, as mentioned above. Of the 39 T7 tRNAs studied in droplets, 22 (56%) were sensitive to this Mg^2+^ concentration range while 17 (44%) were insensitive, as judged by whether the accuracy increased between Conditions A and D (Figure 5E, Table S32). In contrast, only 9 of the 45 NM tRNAs studied in droplets (20%) were sensitive to this Mg^2+^ concentration range while 36 (80%) were insensitive (Table S32). To confirm these observations, we performed UV-detected melts of four T7 tRNAs over a range of Mg^2+^ concentrations (0, 0.5, 2, and 10 mM). We chose two T7 tRNAs that were Mg^2+^-sensitive as both T7 and NM tRNAs (Leu-CAG-1-1, Thr-CGT-1-1) and two T7 tRNAs that were Mg^2+^-insensitive as both T7 and NM tRNAs (Ala-GGC-1-1, Gly-CCC-1-1). We observed that three of the tRNAs behaved as expected. Both Leu-CAG-1-1 and Thr-CGT-1-1 T7 tRNAs had large Tm increases between 0.5 and 10 mM Mg^2+^, while the Tm of the T7 Gly-CCC-1-1 increased only modestly over the same range (Figure S28, Table S33). However, the T7 Ala-GCC-1-1 which appeared to be Mg^2+^-insensitive from the DMS-MaP experiments, was strongly sensitive to Mg^2+^ over this range. Overall, some tRNAs fold with a modest dependence on Mg^2+^ concentration, especially NM tRNAs, while others require higher concentrations of Mg^2+^, which can be present in membraneless compartments (*25*, *26*).

## Discussion

Compartmentalization is thought to have been critical for autocatalytic molecules to arise from the prebiotic milieu (*79, 80*). RNA encapsulation in droplets formed by phase separation is a recently-recognized feature of extant biology, and on the early Earth such compartments could have accumulated otherwise scarce functional RNAs. Complex coacervation is a general property of oppositely charged polyions, but it was unclear whether the resulting compartments would be suitable for native RNA folding and function. The peptide-rich coacervate droplets serving as RNA compartments in this work are very simple homopeptide oligomers lacking biospecific interactions with RNA. Such short chemically simple polymers could have been produced by abiotic processes, for example on mineral surfaces (*16*, *80*), in eutectics (*81*), and as a result of wet-dry cycling (*82*, *83*).

In this work, we demonstrated conditions in droplets that were sufficient to attain native folding of tRNAs, which is essential for RNA function. These coacervate droplets were comprised of lysine and aspartate, both of which are important in extant intrinsically disordered proteins that are known to phase separate either by themselves or with RNA (*84*). Among the conditions that we tested, we found little difference in the physical properties of the droplets, which had appreciable fluidity, even when accounting for changes in the polyion length, and ionic strength. However, despite all conditions having similar half-lives of recovery, apparent diffusion coefficients, and strong RNA accumulation into the coacervate phase, there was a significant impact of the different conditions on the RNA structure.

Using conditions ranging from stringent to mild, which we identified from ILP on a single tRNA, we adapted and applied tRNA structure-seq to study tRNAs in droplets for the first time. We studied ^~^40 different T7 transcripts under several different conditions, which demonstrated that this technique can be applied to multiple RNAs in droplets under multiple conditions. The T7 transcripts of the *E. coli* tRNAs displayed a range of folding ability demonstrating that different sequences have different abilities to fold in the droplets. This appears to be due at least in part to the Mg^2+^ sensitivity of the sequence or to population of misfolded states with near-native stability.

One critical feature of our studies was the ability to directly compare the folding of naturally modified tRNAs to their unmodified T7 transcript counterparts upon encapsulation. Such experiments cannot be performed *in vivo* because tRNAs are always naturally modified. To gain deeper insight into the influence of covalent RNA modifications on native tRNA folding, we purified naturally modified tRNAs from *E. coli* and studied their structures in droplets in the same conditions as the T7 transcripts using tRNA structure-seq. In nearly all cases, the naturally modified versions folded stronger and more natively than the T7 transcripts in the droplets suggesting that these modifications make the RNAs more resistant to unfolding and more active. This was seen with more negative ΔGs of folding and higher A58 reactivates for the naturally modified versions. Unlike the accuracy which has an upper boundary of 100%, A58 reactivity and the apparent ΔGs of folding are not bounded and can go higher or lower depending on the extent of native folding. DMS reactivities at A58 and the 3’-terminal CCA were both high, corresponding to native folding of a reverse Hoogsteen at A58•U54 and extended stacking at the 3’-end of the acceptor stem. The positive impact of covalent RNA modifications on native folding can be seen in the improved prediction accuracy of NM tRNAs over T7 tRNAs.

Special attention was given to the apparent Mg^2+^ sensitivity of the tRNAs. This is of particular importance inside of membraneless compartments, where much of the internal Mg^2+^ may be sequestered in interactions with other polyanions, such as the aspartate carboxylate moieties herein. Again, a potential role of the natural modifications emerges, as most of the naturally modified tRNAs tested appeared to be Mg^2+^ insensitive, perhaps because the natural modifications bias the fold towards the native state. This suggests that high concentrations of Mg^2+^ are not necessary for all tRNAs to fold natively, which may have been advantageous on the early Earth particularly in oceans where Mg^2+^ is far less concentrated than other cations like Na^+^ (*15*).

Natural modifications can enhance RNA folding by lowering the conformational flexibility of the phosphate backbone and reducing the number of possible misfolds. Some important modifications, such as dihydrouridine (*56*, *57*) reduce helix stability, while others like pseudouridine stabilize RNA structure (*55*). Underscoring the importance of natural modifications, loss of tRNA modifications *in vivo* has been implicated in disease (*85*). These diseases are often associated with hypomodification of mitochondrial tRNAs, which can lead to severe phenotypes such as in mitochondrial myopathy encephalopathy lactic acidosis and stroke-like episodes (MELAS); additionally, hypomodification of tRNAs has been implicated in different cancers (*86*). Recent studies have revealed essential roles of covalent RNA modifications in mRNA vaccines (*87*). In summary, this study represents a significant step forward in our understanding of RNA folding in membraneless compartments and the potential of natural modifications to stabilize RNA structures in extant biology, delivery vehicles for RNA therapeutics, and origins of life studies.

## ACKNOWLEDGEMENTS

We thank Professors Squire Booker and Christopher House for their helpful comments and encouragement.

## Authors contributions

M.O.M., C.D.K., and P.C.B. conceptualized the study. M.O.M., R.Y., and S.C. performed the experiments. C.D.K. and P.C.B. acquired the funding. M.O.M., C.D.K., and P.C.B. wrote the original draft. All authors performed reviewing and editing.

## Data and materials availability

All data will be release upon peer review of the manuscript.

## SUPPLEMENTARY MATERIALS

Will be made available upon peer review of the manuscript.

